# (E,E)-bisantrene suppresses *MYC* expression and displays anti-leukemic activity in acute myeloid leukemia

**DOI:** 10.64898/2026.09.10.750805

**Authors:** Heather C. Murray, Kasey Miller, Joshua Brzozowski, Dylan Kiltschewskij, Ishani Roy, Nikita Panicker, Ameha S. Woldu, Benjamin. J. Buckley, Marinella Messina, Chelsea Penney, Peter Cuthbertson, Sumit Sahni, Daniel Tillett, Michael J. Kelso, Jonathan Sillar, Murray Cairns, Anoop Enjeti, Nicole M. Verrills

**Affiliations:** School of Biomedical Sciences and Pharmacy, College of Health, Medicine and Wellbeing, University of Newcastle, Newcastle, New South Wales, Australia; Precision Medicine Program, Hunter Medical Research Institute, New Lambton, New South Wales, Australia; Racura Oncology Limited, Sydney, New South Wales, Australia; Calvary Mater Newcastle Hospital, Waratah, NSW, Australia; School of Medicine and Public Health, The University of Newcastle, Callaghan, NSW, Australia

## Abstract

**Background:** Acute myeloid leukemia (AML) is genetically diverse with a high unmet clinical need for improved treatment options. Dysregulation of the transcription factor MYC plays a central role in AML progression and therapeutic resistance. (E,E)-bisantrene was recently found to inhibit *MYC* transcription and downstream activity via G-quadruplex DNA stabilization. This study aimed to evaluate the mechanism of action and preclinical activity of (E,E)-bisantrene in AML.

**Methods:** The *in vitro* and *in vivo* activity of (E,E)-bisantrene was determined in a variety of AML models (cell lines, xenograft mouse models, and *ex vivo* human AML mononuclear cells). Transcriptomic, proteomic and phosphoproteomic analyses were performed after treatment with (E,E)-bisantrene. Analyses of -omics data to identify enriched pathways and upstream regulators were performed.

**Results:** (E,E)-bisantrene demonstrated potent anti-proliferative activity across a panel of AML cell lines, inducing apoptosis and reducing S phase proportions. (E,E)-bisantrene significantly prolonged survival in cell- and patient-derived xenograft models of AML. Mechanistically, RNA-seq and proteomic analysis of MOLM13 and MV4-11 cells treated with (E,E)-bisantrene showed significant reductions in the activity of MYC and E2F, together with the cell cycle regulators CDK1/2/4/5. Transcript and protein levels of *MYC* were reduced in a dose- and time-dependent manner. TP53 and inflammation-associated transcript signatures were also observed.

**Conclusion:** Anti-proliferative activity of (E,E)-bisantrene in preclinical AML models was associated with a downregulation of MYC, CDK1/2/4/5 and E2F. This study supports the ongoing clinical evaluation of (E,E)-bisantrene in AML where MYC is a clinically relevant driver of disease aggressiveness and therapy resistance.

## 1. Introduction

Acute myeloid leukemia (AML) is a genetically diverse malignancy characterized by abnormal growth and differentiation of hematopoietic stem cells, leading to accumulation of immature myeloid precursor cells in the bone marrow and peripheral blood.^1^ Most transplant-eligible AML patients receive first line combination therapy of daunorubicin and cytarabine for induction before stem cell transplation.^1^ Although this treatment is effective in achieving remission in ∼60% patients, the majority relapse within 5 years.^2, 3^ Long term outcomes in AML remain dismal, with a 5-year survival rate of only 35%.^4^ The severe toxicity associated with high intensity induction chemotherapy is a major clinical challenge and precludes many patients from receiving effective treatments.^5,6^

The recent introduction of novel targeted therapies for AML treatment has provided less toxic alternatives for AML patients.^7^ Unfortunately, the genetic diversity of AML means targeted therapies are effective in only relative small proportions of patients, with many patients having malignancies with no actionable driver mutations.^8^ Moreover, even in patients that respond to targeted therapy, most responders eventually acquire resistance and relapse.^9^ Novel treatment approaches of broad utility are urgently needed.

MYC is a central oncogenic transcription factor implicated in cellular proliferation, metabolic regulation, stemness, differentiation blockade, and therapeutic resistance across multiple cancers.^10^ MYC is overexpressed in up to 90% of AML malignancies and has been associated with aggressive disease biology and adverse prognostic outcomes.^11^ The central role of MYC in AML makes it an attractive target for drug development, however, the intrinsically disordered protein structure of MYC has made targeting challenging using conventional small molecule targeted approaches.^12^

Stabilization of the G-quadruplex (G4) DNA structure within the *MYC* gene promoter region is a promising alternative strategy for suppressing MYC activity.^12^ In this approach, silencing of *MYC* transcription through small molecule binding to a G4 structure contained within the *MYC* promoter results in a reduction in MYC protein levels and downstream MYC-driven oncogenic pathways important for cancer cell survival and proliferation.

(E,E)-bisantrene has been shown to suppress *MYC* expression in cancer cells, including AML.^13^ Recent studies found that only the (E,E)-isomer of bisantrene is able to inhibit *MYC* transcription via binding to and stabilizing the *MYC* promotor G4 region.^14^

In addition to the transcriptional silencing of *MYC*, bisantrene elicits a range of additional anti- cancer activities including: (i) telomerase inhibition; (ii) topoisomerase-IIα inhibition; (iii) DNA and RNA synthesis inhibition; (iv) immune-stimulation; (v) angiogenesis inhibition, and (vi) increasing m^6^A RNA methylation levels.^13, 15–24^ These multiple complementary mechanisms of action may contribute to the broad anti-leukemic activity of bisantrene across genetically heterogeneous AML populations, and may reduce therapy resistance arising compared to agents targeting a single oncogenic pathway. This broad anti-leukemic activity was explored in two recent phase 2 clinical trials which found (E,E)-bisantrene is an effective salvage therapy for relapsed and refractory AML (R/R AML) independent of underlying mutational status.^25, 26^

In this study, the anti-cancer effects and mechanisms of action of (E,E)-bisantrene was investigated in a range of preclinical AML models. (E,E)-bisantrene exhibited robust anti-leukemic activity, both *in vitro* and *in vivo*. Transcriptomic, proteomic and phosphoproteomic analyses identified *MYC* gene expression silencing as a key driver of (E,E)-bisantrene activity in AML.

## 2. Materials and Methods

### 2.1. Cell Lines and Drugs

Human AML cell lines THP1, MOLM13, MV4-11, and HL60 were sourced as described previously.^27^ OCI-AML3 was a kind gift from Heather Lee (The University of Newcastle, Australia). M07e was a kind gift from Leonie Ashman (The University of Newcastle, Australia). Kasumi1 was purchased from the American Type Culture Collection (ATCC; Manassas). U937 and MonoMac6 were purchased from AcceGen (Fairfield, New Jersey). All lines were cultured in a humified chamber at 37 °C with 5% CO_2_ in RPMI supplemented with 10% fetal bovine serum (FBS), 20 mM HEPES and 2 mM L-glutamine. THP1 cells were additionally supplemented with 50 µM β-mercaptoethanol. A panel of human AML cell lines (NB4, MV4-11, MOLM13, AP1060, HL60, PL21, NOMO1, THP1, K-562, Kasumi1, SET2, HEL, KG1) was tested independently by Oncolines B.V. (Oss, Netherlands) according to their standard operating procedures.

Mouse myeloid early progenitor FDCP1 cell lines transduced with either empty vector (EV) or oncogenic mutant AML-associated mutations (*KIT-D816V, FLT3-ITD, KRAS-G12V*) were generated as previously described.^28, 29^ The human *KRAS G12V* vector was a kind gift from Dr Chen Chen Jiang (University of Newcastle, Australia). FDCP1 lines were cultured in a humified chamber at 37 °C with 5% CO_2_ in DMEM supplemented with 10% FBS, 2 mM L-glutamine, and 20 mM HEPES. FDCP1 EV cells were additionally supplemented with 0.5 ng/mL granulocyte-macrophage colony-stimulating factor (GM-CSF). Media for the KIT-D816V cell line was supplemented with 1 mg/mL G418 for one week after thawing and then removed.

AML patients were recruited through the Calvary Mater Newcastle (Australia) in accordance with institutional guidelines. Studies were approved by the human ethics committees of the Hunter New England Area Health (HNEAH) service (2019/ETH00744) and the University of Newcastle (H-2011-0062). Written informed consent was obtained from all participants. Karyotype and *FLT3* status were determined through routine pathology tests. Presence of other AML-associated mutations were identified through routine clinical tests using next generation sequencing. Mononuclear cells were isolated from bone marrow samples using Lymphoprep density gradient medium (StemCell; Vancouver, Canada) and SepMate tubes (StemCell), as previously described.^27^ A blast count cut-off of ≥30% in the bone marrow or peripheral blood was used for selection of primary AML samples.

AML cells were isolated from the spleens of immunocompromised mice engrafted with AML PDXs 5, 16, or 18.^30^ Spleens were passed through a 40 µm filter, followed by red blood cell lysis to remove splenocytes. AML-associated mutations were retrieved from Bruedigam et al.^31^

(E,E)-bisantrene dihydrochloride ((E,E)-bisantrene; Racura Oncology Ltd, Australia) was prepared as 20 or 50 mM stocks in DMSO, aliquoted and stored at -20 °C to reduce freeze-thaw cycles. Care was taken to prevent exposure to light for all solutions containing (E,E)-bisantrene, to minimize photoisomerization to (E,Z)-bisantrene.

### 2.2. Cell Viability Assay

Cell viability was determined using a resazurin metabolic activity assay, as described previously.^28^ For cell viability studies, cells were plated out in duplicate wells of 96-well microtiter plates at cell densities depending on their growth rates; 2x10^4^ cells/well (MOLM13), 4x10^4^ cells/well (MV4-11, THP1), 6x10^4^ cells/well (M07e), 10x10^4^ cells/well (OCI-AML3), 8x10^4^ cells/well (HL60), 1.6x10^5^ cells/well (Kasumi1), 2x10^4^ cells/well (MonoMac6), 2x10^4^ cells/well (U937), or 1x10^4^ cells/well (all FDCP1 lines). Drug was then added and cells incubated for 72 h. At the end of the treatment, viability was determined by the resazurin assay.^27^ Graphpad Prism software (La Jolla, CA, USA) was used to generate graphs and IC_50_ values. Viability was normalized to untreated cells at 100%.

The Oncolines AML cell line panel was tested for cell viability after 72 h (E,E)-bisantrene treatment using the ATPlite assay (Revvity, Waltham, MA, USA).

### 2.3. Primary AML Ex vivo Assay

The ATPlite assay for primary AML samples and PDX cells *ex vivo* was performed in 96-well plates with 5x10^4^ cells/well (5x10^5^ cells/ml) or 384-well plates with 1x10^4^ cells/well (5x10^5^ cells/ml) in IMDM supplemented with 0.5% FBS. (E,E)-bisantrene (1 µM) was added and cell viability determined by ATPlite assay after incubation for 24 h at 37 °C, 5% CO_2_ according to the manufacturer’s instructions (Revvity). Cell viability was normalized to untreated cells at 100%.

### 2.4. Apoptosis Assay

Apoptosis was assessed by Annexin V-APC flow cytometry assays (BD Biosciences) according to the manufacturer’s instructions. Samples were analyzed on a Canto II flow cytometer and data analyzed using Flow Jo (BD Biosciences).

### 2.5. Cell Cycle Assessment

Cell cycle was assessed by propidium iodide staining and flow cytometry, as described previously.^27^ Data analysis was performed using the Watson Pragmatic model in FlowJo (BD Biosciences). To ensure consistent modeling of S-phase and G2/M populations across samples, the coefficient of variation (CV) of the G2 peak was constrained to be equal to the CV of the G1 peak. Data was normalized to 100% for each replicate.

### 2.6. Mouse Models

#### 2.6.1. AML Cell Line Derived Xenograft (CDX) Model

Mouse procedures were performed in accordance with institutional approvals (A-2020-010). Mouse xenograft models utilized NOD.Cg-Prkdcscid Il2rgtm1Wjl/SzJ (NSG) mice. Luciferase tagged MOLM13 (MOLM13-luc) cells were a kind gift from Charles de Bock (Children’s Cancer Institute, Australia). Mice were xenografted with MOLM13-luc (5x10^5^ cells/animal) via tail vein injection. Leukemia burden throughout the study was monitored by bioluminescence imaging (BLI). BLI was first performed between days 5 and 7 post-engraftment; animals with detectable BLI readings were selected and randomized into groups for treatment. Mice were treated with vehicle control or 5 mg/kg *i.v.* or 10 mg/kg *i.p* (E,E)-bisantrene 2-3 times/week (total of 8 doses) over 3 weeks. (E,E)-bisantrene injection solutions were prepared by dissolving the dihydrochloride salt (CAS No. 78186-34-2; Racura Oncology Ltd, Australia) in 8% Captisol (sulfobutylether β-cyclodextrin; CyDex Pharmaceuticals) in water.

Following commencement of treatment, BLI was conducted weekly. Mice were injected *i.p.* with 3 mg D-luciferin (Promega #P1043; 150 mg/kg in PBS), anesthetized using 2-5% isoflurane (500-1000 CC/min) and imaged approximately 10 minutes after D-luciferin injection using a Xenogen IVIS Spectrum imager (Revvity). Images were analyzed using Living Image software (Revvity). A rectangular region of interest (ROI) was placed over each individual mouse and total flux (p/s) within each ROI was calculated. Mice were monitored until ethical endpoint.

#### 2.6.2. AML Patient Derived Xenograft (PDX) Model

NSG mice were engrafted with 2x10^6^ PDX-AML16 cells via tail vein injection.^30^ Leukemia burden was determined weekly by collection of peripheral blood from the tail vein and flow cytometry analysis to determine the proportion of hCD45^+^/(hCD45^+^ and mCD45^+^) cells. Whole blood was subjected to red blood cell lysis and fixation prior to staining with BV421-conjugated human CD45 (hCD45) and APC-Cy7-conjugated mouse CD45 (mCD45) antibodies (BD Biosciences). Stained cells were analyzed using a FACSCanto II flow cytometer (BD Biosciences) and the percentage of hCD45^+^ cells was calculated. Once mice achieved detectable levels of hCD45^+^ cells in the peripheral blood (approximately 3.5 weeks following engraftment), mice were randomized into groups and treated with vehicle control or (E,E)-bisantrene (2.5 or 5 mg/kg; *i.v.;* 3 times/week). Following randomization, blood was collected twice weekly from the tail vein, typically before (E,E)-bisantrene treatment. Drug was administered in two cycles; each cycle consisted of two weeks of treatment with a drug free period of 12 days between cycles. The ethical endpoint was reached once hCD45^+^ cells reached 25% in peripheral blood.^32^

### 2.7. RNA-sequencing Analysis

RNA-seq was conducted using the MAZTER-Seq method to enable quantification of both mRNA expression and putative m^6^A events (for a separate study).^33^ For each sample, 5 µg total RNA was subject to poly(A) enrichment utilizing Dynabeads (Thermo Fisher) as per the manufacturer’s instructions. Samples were then treated with 10U MazF endoribonuclease (TakaraBio), which selectively cleaves RNA at unmethylated ACA motifs, followed by end repair using 4 µL T4 PNK (New England Biolabs). Library preparation was subsequently performed using a SMARTer smRNA-Seq Kit for Illumina (TakaraBio); this kit was selected to preserve sequence information at the 5′ and 3′ ends for future analysis of ACA cleavage. Briefly, cleaved RNA was polyadenylated in the presence of ATP, primed with SMARTer 3′ smRNA dT primers, and reverse transcribed using 2 µL PrimerScript reverse transcriptase (200U/µL). cDNA was amplified via 10 PCR cycles using uniquely indexed PCR primers and validated by High Sensitivity DNA Bioanalyzer Kit (Agilent). Samples were then normalized to 0.5 ng/µL, pooled and subjected to 151 cycles of paired end sequencing using an Illumina NovaSeq 6000 instrument with S2 flow cell and 300-600 pM loading concentration.

Raw bcl files were demultiplexed into fastq format using *bcl2fastq*, and lane data were merged for each sample. Initial quality control was performed using *fastp* (v.0.20.0),^34^ removing: the first three 5′ and 3′ bases (as recommended by the SMARTer smRNA-Seq Kit for Illumina manual), 3′ bases with Phred quality score < 28, and fragments ≤ 25 nt. Fastq files were then aligned to the reference genome (GRCh38, NCBI) using *star* (v.2.5.3a),^35^ and reads aligning to genes were counted using *HTSeq* (v.0.12.4).^36^ Next, raw counts were merged into a single matrix, genes with low counts (counts-per million equal to ≤ 5 raw counts in the smallest library) in > 50% samples were removed and clustering of biological replicates was examined via multidimensional scaling. The *EdgeR* library (v.3.42.4) was then used to calculate normalization factors, estimate dispersion, and analyze pairwise differential gene expression between treated and untreated cells. Correction for multiple testing was performed using a Benjamini-Hochberg false discovery rate.

To interrogate the expression of genes known to interact with MYC and TP53, protein-protein interaction networks were constructed for both genes using STRINGdb ^37^. All included nodes required a minimum interaction score of 0.9 (“highest confidence”) and supported by both experimental data and existing interaction databases. Each network was further restricted to the top 20 nodes (ranked by interaction score) to focus on the most biologically salient proteins.

The RNAseq data has been uploaded to the Gene Expression Omnibus, accession number GSE346849.

### 2.8. Proteomic and Phosphoproteomic Analysis

Label free proteomic and phosphoproteomic profiling was performed using the EasyPhos method, as previously described.^38, 39^ Peptides were preconcentrated using an EASY-Spray PepMap C18 75 μm × 20 mm column (Thermo Fisher Scientific). Separation was achieved using a 75 μm ✕ 25 cm EASY-Spray PepMap C18 column (Thermo Fisher Scientific) on an Orbitrap Exploris 480 (Thermo Fisher Scientific); proteome: 5–35% solvent B over 90 min; and phosphoproteome: 5–30% solvent B over 90 min (solvent B: 80% acetonitrile, 0.1% formic acid). Samples were analyzed using data dependent acquisition, with full MS scans of 360-1500 *m/z* acquired at a resolution of 60,000, normalized automatic gain control of 300% and maximum injection time of 100 ms. MS/MS fragments were measured at a resolution of 15,000, automatic gain control of 100%, a normalized collision energy of 30 and 36, and automatic maximum injection time.

Raw files were analyzed using Proteome Discoverer 2.5 (Thermo Fisher Scientific), as previously described.^38^ Differential expression (|fold change| >1.5) between treatment groups was assessed using Student’s *t* tests, and p<0.05 was considered significant. Only entries with values in all replicates of at least one sample group were considered significant.

The mass spectrometry proteomics data have been deposited with the ProteomeXchange Consortium (http://proteomecentral.proteomexchange.org) via the PRIDE partner repository^40^ using the dataset identifier identifier PXD083214 and DOI:10.6019/PXD083214.

### 2.9 DepMap

Transcriptomic data for MV4-11, M07e, MOLM13, U937 (more sensitive to (E,E)-bisantrene), and THP1, OCIAML3, MonoMac6, HL60, and Kasumi1 (less sensitive to (E,E)-bisantrene) was downloaded from depmap.org (DepMap, Broad (2026). DepMap Expression Public 26Q1 dataset).^41^

### 2.10 Pathway and Kinase Enrichment Analysis

Upstream regulators of differentially expressed genes (q<0.05) or proteins (p<0.05, without a fold change or number of values cut off) were identified using Ingenuity Pathway Analysis (IPA) software. Upstream regulators which were predicted to be activated (z score ≥ 2, p<0.05) or inhibited (z score ≤ -2, p<0.05) were identified. Pathway enrichment was assessed using Gene Set Enrichment Analysis software (GSEA; Broad Institute)^42, 43^ and the Hallmark MsigDB database. Kinase enrichment analysis was performed using the phosphoproteomic datasets. KSEApp^44^ was used to identify enrichment within curated (PhosphositePlus^45^) kinase-substrate links. PTM-SEA^46^ was used as a complementary kinase enrichment analysis evaluation, using the human ptmsigdb v2.0 database.

### 2.11 Western blotting

Western blotting of whole cell lysates was performed using standard procedures, as previously described^27^. Primary antibodies used were MYC (ab32072, Abcam, 1:1000), P53 (SC-126, Santa Cruz Biotechnology, 1:1000), p21 (2947S, Cell Signaling Technology (CST), 1:1000), MYC (CST #5605S 1:500), GAPDH (CST #97166S, 1:5000), and species-specific HRP- (CST #7074S and #7076S) or LI-COR IR-dye conjugated (926-32213, 926-32212) secondary antibodies. Membranes were imaged using an Amersham 600 gel imager (GE Healthcare) or a LI-COR Odyssey M infrared imaging system. Band intensities were quantified using ImageJ v1.54p or Image Studio software (LI-COR). Rhodamine-conjugated GAPDH (BioRad (12004167)) loading controls were imaged using a ChemiDoc (BioRad) and analyzed using Image Lab software (BioRad). Results were normalized to the loading control and expressed as a fold-change relative to control (untreated). Data represent at least three independent biological replicates and are presented as mean ± SEM. Statistical significance was determined by one-way ANOVA with posthoc Tukey HSD test compared to the untreated control, with p<0.05.

### 2.12 qPCR

Total RNA was extracted from cell pellets using TRIreagent (Merck, Cat# T9424) or Bioline ISOLATE II Mini Kit (Meridian Science) according to the manufacturer’s instructions. RNA concentrations were determined using a NanoDrop^®^ spectrophotometer. Equal amounts of RNA were reverse-transcribed into complementary DNA (cDNA) using LunaScript RT SuperMix Kit (New England Biolabs, Cat# M3010L) or qScript IITM cDNA Supermix Kit with random hexamers or oligo(dT) primers (Quantabio). qPCR reactions were prepared using either TaqMan Gene Expression Assay (Thermo Fisher Scientific; *MYC*: Cat# HS00153408_m1; *GAPDH*: Cat# HS02786624_g1) or using PowerUp SYBR Green master mix (Thermo Fisher Scientific), gene-specific primers,^13^ and diluted cDNA template. Samples were analyzed using the QuantStudio™ 3 or 5 PCR System (Thermo Fisher) under default cycling conditions, including an initial denaturation step followed by 40 cycles of amplification. Melt curve analysis was performed for SYBR Green assays to confirm amplification specificity. Gene expression levels were calculated using the comparative Ct (2⁻^ΔΔCt^) method and normalized to housekeeping gene (*GAPDH*). All reactions were performed in technical triplicates with at least three biological replicates. Statistical significance was determined by one-way ANOVA with posthoc Tukey HSD test compared to the untreated control, with *p* < 0.05.

### 2.13 Data Analysis

Statistical comparisons were performed using GraphPad Prism or Microsoft Excel. Survival was analyzed using Kaplan Meier curves and compared by log-rank test. Apoptosis and cell cycle assays were analysed by 2-ANOVA with Fishers LSD. Annotation of RNAseq, proteomic, and phosphoproteomic data was performed with the assistance of StringDB^37^ and Venny^47^.

## 3 Results

### 3.1 Effect of (E,E)-bisantrene in human AML cell lines

The effect of (E,E)-bisantrene on cell viability was determined using the resazurin metabolic assay in a panel of human AML cell lines. Most cell lines were sensitive to nanomolar concentrations of (E,E)-bisantrene (**Figure 1A**). M07e cells were the most sensitive (IC_50_: 14.1 nM), followed by MV4-11 (IC_50_: 15.0 nM) and MOLM13 (IC_50_: 34.8 nM) cells, while Kasumi1 cells were the least sensitive (IC_50_: >200 nM). As bisantrene is a known substrate for the drug efflux pump P-gp,^48^ reduced sensitivity to Kasumi1 cells could be due to high expression of P-gp in these cells (**Table S1**).

**Figure 1:**
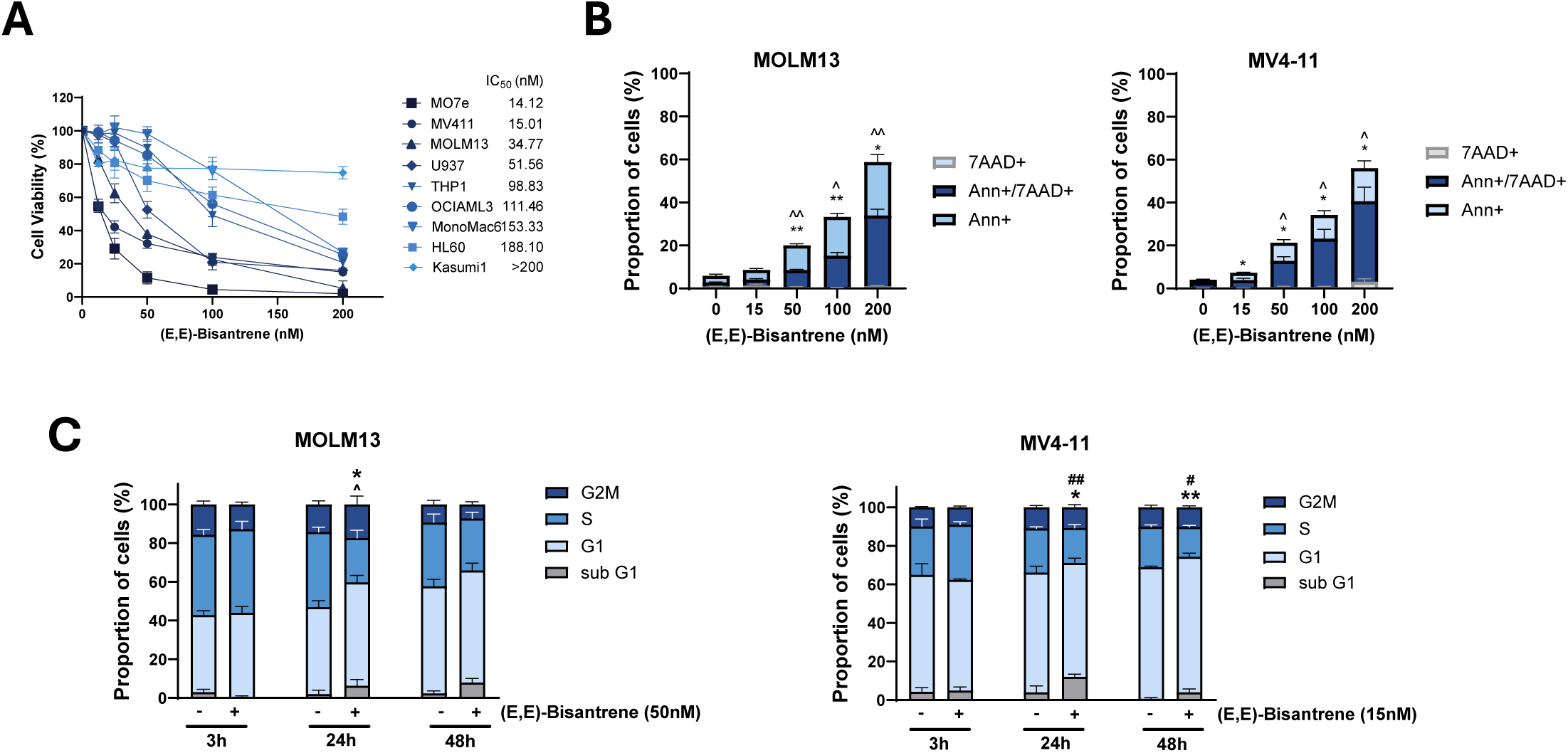
Anti-cancer activity of (E,E)-bisantrene in AML cell lines. **(A)** 72 h dose-response curves of human AML cell lines. Inset: IC_50_ was determined for each cell line, n=3. **(B)** Apoptosis was assessed using Annexin V/7AAD assays at 48 h in MOLM13 and MV4-11 human AML cell lines, n=3, mean ± SEM, *p<0.05, **p<0.01 Ann+; ^p<0.05, ^^p<0.01 Ann+/7AAD+ compared to untreated cells ((E,E)-bisantrene (0 nM)). **(C)** Cell cycle response to (E,E)-bisantrene was measured at 3, 24, and 48 h in MOLM13 and MV4-11 human AML cell lines, n=3. Mean +/-SEM. Difference in S-phase proportion between (E,E)-bisantrene treated and untreated cells (*p<0.05; **p<0.01); difference in G1-phase proportion (^p<0.05); and sub-G1 phase (^#^p<0.05; ^##^p<0.01) in (E,E)-bisantrene treated cells and, 2-way ANOVA.

Independent assessment of the anti-proliferative activity of (E,E)-bisantrene in the Oncolines AML cell line panel using the ATPlite assay showed similarly high (E,E)-bisantrene sensitivity in most cell lines (9 of 13 showed IC_50_ < 200 nM) (**Table S1**). Consistent with the resazurin assay, MV4-11 (IC_50_: 26 nM) and MOLM13 (IC_50_: 32 nM) showed high sensitivity to (E,E)-bisantrene, while Kasumi1 was less sensitive (IC_50_: 206 nM). Cell lines with reduced sensitivity to (E,E)-bisantrene (Kasumi1, SET2, HEL and KG1) all showed high P-gp expression (**Table S1**).

### 3.2 Effect of (E,E)-bisantrene in mouse myeloid progenitor cells

AML is commonly driven by mutations in oncogenes (e.g., *FLT3, KIT, KRAS*) that can affect drug sensitivity. Two of the human AML cell lines that were highly sensitive to (E,E)-bisantrene, MV4- 11 and MOLM13, are positive for the *FLT3* internal tandem duplication (*FLT3-ITD*), while the mutant *KIT* (*KIT-N822K*) AML cell line Kasumi1 was the least sensitive. To directly investigate the relationship between AML associated oncogenes and (E,E)-bisantrene sensitivity, we utilized isogenic FDCP1 mouse myeloid progenitor cells, which are dependent on GM-CSF or IL3 for survival. Transduction of these cells with oncogenic mutant *FLT3-ITD*, *KIT-D816V*, or *KRAS- G12V* results in factor-independent growth, and *FLT3-ITD* and *KIT-D816V* induce leukemogenesis *in-vivo.*^28, 49^ The empty vector (EV) cells, transduced with an empty vector pRUFneo retroviral construct and grown in GM-CSF, were used as a control. All cells were analyzed for (E,E)-bisantrene sensitivity using the 72 h resazurin assay.

The EV and transduced lines were all highly sensitive to (E,E)-bisantrene, with IC_50_ values in the low nanomolar range (8-16 nM; **Supplementary Figure S1**), indicating that (E,E)- bisantrene is potent against rapidly proliferating early progenitor myeloid cells *in vitro* irrespective of whether they use cytokine receptor signaling or oncogenic mutant kinase activated signaling for survival.

### 3.3 Effect of (E,E)-bisantrene on apoptosis and cell cycle

MOLM13 and MV4-11 cells were selected for further analysis due to their high sensitivity to (E,E)-bisantrene (**Figure 1A, Table S1**). To determine if inhibition of cell viability in response to (E,E)-bisantrene was due to the induction of apoptosis, MOLM13 and MV4-11 cells were treated with a dose range of (E,E)-bisantrene (15, 50, 100, 200 nM) for 48 hours. A dose-dependent increase in the proportion of cells undergoing early (Annexin+/7AAD-) and late (Annexin+/7AAD+) stage apoptosis was observed in both cell lines (**Figure 1B**).

Next, the effect of (E,E)-bisantrene on cell cycle was assessed using flow cytometry, using 50 nM (MOLM13) and 15 nM (MV4-11) (E,E)-bisantrene concentrations, to approximate the respective IC_50_ values. In MOLM13 cells the proportion of cells in G1 phase was increased, and cells in S phase was reduced after 24 h incubation with (E,E)-bisantrene, respectively (**Figure 1C**) (E,E,)-bisantrene also reduced the S-phase proportion and increased sub-G1 in the MV4-11 cells at 24 and 48 h (**Figure 1C**). Collectively, these studies show that treatment with (E,E)-bisantrene results in G1/S phase cell cycle arrest and induction of apoptosis.

### 3.4 Anti-cancer activity of (E,E)-bisantrene against primary human AML blasts

To assess the activity of (E,E)-bisantrene on human primary AML blasts, bone marrow samples were collected from AML patients (n=25) and mononuclear cells isolated. The *ex vivo* sensitivity of three AML patient derived xenograft (PDX) samples, expanded *in vivo* and isolated from spleen, was also assessed (n=3). The effect of 1 µM (E,E)-bisantrene on blast viability at 24 h was determined using an ATPlite assay. A range of sensitivities was observed, with 5/25 patient samples displaying a normalized viability of ≤ 50% (**Figure 2**). AML samples with *NPMI* mutation were associated with better *ex vivo* (E,E)-bisantrene response than *NPM1* wildtype AML (p=0.0027, *t-* test), while samples with *IDH2* mutations were associated with lower *ex vivo* (E,E)-bisantrene sensitivity (p=0.049, *t*-test).

**Figure 2:**
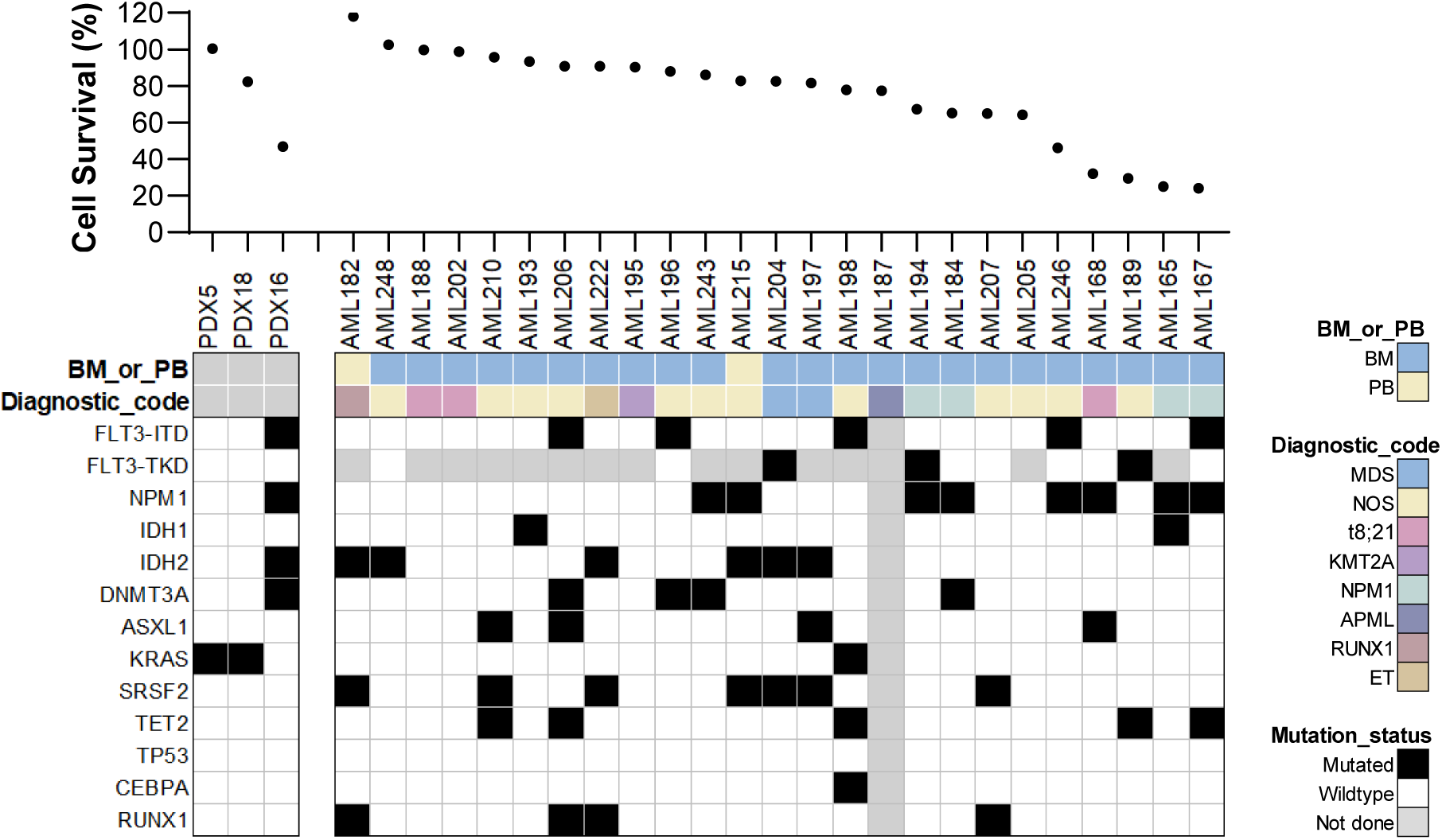
*Ex vivo* response of human AML patient samples treated with (E,E)-bisantrene. Primary human AML blasts, or cells isolated from AML PDXs, were treated with 1 µM (E,E)-bisantrene *ex vivo* for 24 h. Cell viability was measured using ATPlite assay. BM, bone marrow; PB, peripheral blood. Diagnostic code: MDS, AML with myelodysplasia-related changes; NOS, AML not otherwise specified; t(8;21), AML with t(8;21) *RUNX1-RUNX1T1*; KMT2A, AML with t(9;11)(p22;q23); NPM1, AML with mutated *NPM1*; APML, acute promyelocytic leukemia; RUNX1, AML with mutated *RUNX1*; ET, AML transformed from essential thrombocytosis.

### 3.5 (E,E)-bisantrene shows dose dependent anti-leukemic efficacy in AML xenograft models

*In vivo* efficacy of (E,E)-bisantrene was assessed in a cell line xenograft (CDX) model, where MOLM13-luc engrafted NSG mice were treated with (E,E)-bisantrene (5 mg/kg *i.v.* or 10 mg/kg *i.p.*; 2-3 times a week). By week 3, only one vehicle mouse remained, precluding statistical comparison of treated mice to vehicle after 3 weeks. Mice treated with 10 mg/kg (E,E)-bisantrene showed lower bioluminescence than the 5 mg/kg (E,E)-bisantrene treated group (*p=*0.0151) at the end of week 3, indicating a dose dependent effect (**Figure 3A)**. A significant increase in overall survival was seen in mice treated with (E,E)-bisantrene (median survival 20 days (5 mg/kg; p=0.0112) or 24 days (10 mg/kg; p=0.0006)) compared to vehicle treated mice (median survival 17 days; **Figure 3B**).

**Figure 3:**
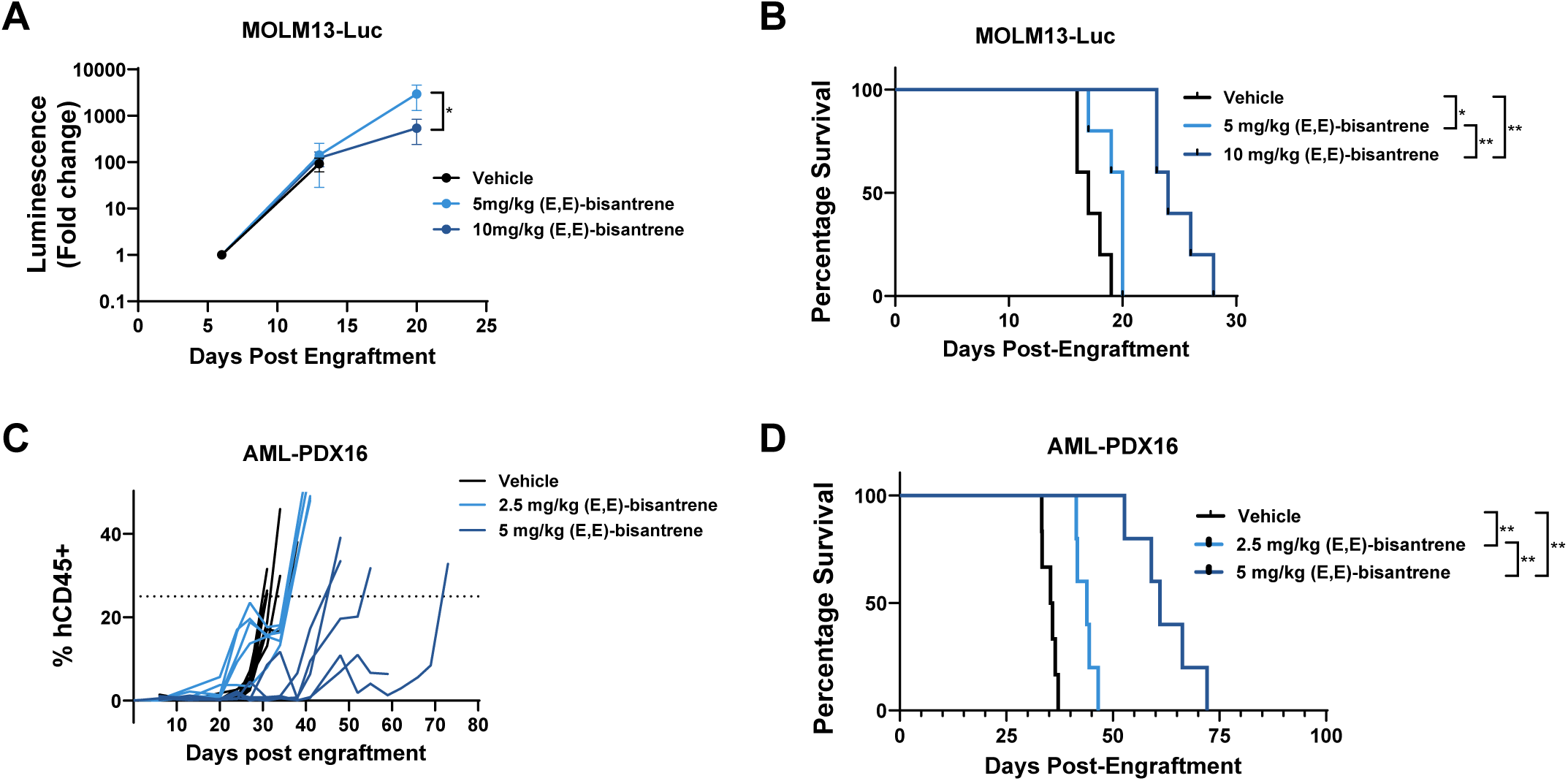
*In vivo* activity of (E,E)-bisantrene in mouse models of AML. **(A,B)** NSG mice were engrafted with MOLM13-luc cells and (E,E)-bisantrene was administered three times per week at 5 mg/kg (*i.v.*) or 10 mg/kg (*i.p.*) for up to 3 weeks. **(A)** Leukemia burden monitored by bioluminescence imaging. *\*p<0.05*, *t*-test. **(B)** Kaplan-Meier survival analysis of MOLM13-luc xenografts. N≥5 mice/group. Median survival: vehicle = 17 days, 5 mg/kg (E,E)-bisantrene = 20 days, 10 mg/kg (E,E)-bisantrene = 24 days. **(C,D)** AML-PDX16 cells were engrafted into NSG mice and once human CD45+ cells reached ∼1% in the peripheral blood, mice were randomized and treated with vehicle control, 2.5 mg/kg or 5 mg/kg (E,E)-bisantrene (*i.v.*) 3 times a week. **(C)** Peripheral blood human CD45 levels (hCD45+), as a percentage of total CD45 positive cells. Mice were sacrificed when the percentage of human CD45+ cells in the peripheral blood reached 25%. **(D)** Kaplan-Meier survival curve. Median survival: vehicle = 35.6 days, 2.5 mg/kg (E,E)-bisantrene = 43.9 days, 5 mg/kg (E,E)-bisantrene = 61 days. n≥5 mice/group. *\*p<0.05, **p<0.01*.

*In vivo* efficacy of (E,E)-bisantrene was further examined in a PDX model of AML (AML16, **Figure 3C,D**). This PDX, which was originally isolated from a 61 year old female with AML M4 subtype, with normal cytogenetics and positive for *FLT3-ITD* and mutant *IDH2*, *NPM1* and *WT1,*^30^ was sensitive to (E,E)-bisantrene in the *ex vivo* viability assay (**Figure 2**). Two doses of (E,E)-bisantrene (2.5 and 5 mg/kg; *i.v.*) administered three times per week were tested in NSG mice engrafted with PDX-AML16 cells. Leukemic burden (% human CD45+ cells) increased rapidly for vehicle control mice, with a median survival of 35.6 days (**Figure 3C,D**). Treatment with 2.5 mg/kg (E,E)-bisantrene had a modest effect on leukemic burden and increased median survival (43.9 days; p=0.0014) compared to vehicle control (**Figure 3C,D**). Mice treated with 5 mg/kg (E,E)-bisantrene showed a greater response to treatment, with a slower increase in leukemic burden compared to vehicle (**Figure 3C**). Survival for the 5 mg/kg (E,E)-bisantrene treated group (median survival: 61 days) was significantly longer than the vehicle control group (p=0.0014) (**Figure 3D**). Overall, (E,E)-bisantrene demonstrated dose dependent anti-leukemic effects in both AML cell- and patient-derived xenograft mouse models.

### 3.6 RNA-seq and proteomic analysis reveals suppression of MYC signaling, and increased inflammatory signaling in response to (E,E)-bisantrene

To investigate the mechanisms of AML action for (E,E)-bisantrene, RNA-seq analysis was performed on MOLM13 and MV4-11 cells treated with 50 nM or 15 nM (E,E)-bisantrene, respectively, for 24 h. Levels of >12,000 genes were quantified in both cell lines. In MOLM13, 2,510 genes were significantly increased, and 2,360 significantly decreased after (E,E)-bisantrene treatment compared to untreated cells (**Figure S2A**, **Table S2**). Top increased transcripts included those for chemokine *CXCL1*, receptor tyrosine kinase *ERBB2*, and HLA antigens *HLA-DRB5*, *HLA-DMB*, and *HLA-DQB1* (**Figure S2A**). Decreased transcripts included those for G-protein signaling regulator *RGS16*, inhibin *INHBE*, and enzyme *PHGDH* (**Figure S2A**). In MV4-11, 1,177 genes were significantly increased, and 1,223 significantly decreased after (E,E)-bisantrene treatment (**Figure S2B**, **Table S2**). Top increased transcripts included those for cell transcription factor *TFAP2B*, contactin-associated *CNTNAP2*, immunoglobulin receptor *FCER1A*, and the CDK inhibitor *CDKN1A* (**Figure S2B**). Decreased transcripts included those for the dehydrogenase/reductase *DHRS2*, tyrosinase-related protein *TYRP1*, and melanocyte protein *PMEL* (**Figure S2B**).

Across the two cells lines, 317 significantly increased (fold change >1.5, q value <0.05) and 385 significantly decreased (fold change <-1.5, q value <0.05) transcripts were shared (**Table S2**). Shared increased transcripts included several for antigen presentation (*HLA-DRB5, HL- DRB1, HLA-DPA1, HLA-DPB1, HLA-DQB1, HLA-DMA, HLA-DMB, CD74*), p53 signaling (*CDKN1A, MDM2, GADD45A*), and several extracellular matrix associated transcripts (*COL9A2*, C*OL4A2, COL6A2, COL6A1, MMP19, MMP25, TIMP2*; **Table S2**). Shared decreased transcripts were related to regulation of cell cycle (*WEE1, E2F1, E2F2, RAD21, CDK2*), cholesterol metabolism (*HMGCR, HMGCS1, MVD, MVK*), and homology directed DNA repair (*RAD1, BRIP1, POLE2, RPA3, BLM*) related transcripts (**Table S2**).

To analyze the effect of (E,E)-bisantrene treatment on the AML proteome, proteomic analysis of MOLM13 and MV4-11 cells treated with (E,E)-bisantrene (MOLM13: 50 nM; MV4- 11: 15 nM) for 24 h was performed (**Tables S3** and **S4, Figure S3**). Few total proteome changes were observed, with 16 and 9 proteins significantly up-regulated, and 15 and 11 proteins significantly down-regulated, in MOLM13 and MV4-11 cells, respectively (**Tables S3** and **S4, Figure S3**). No common differentially expressed proteins were evident between the two cell lines. To further explore the functional implications of the (E,E)-bisantrene induced transcriptome and proteome changes, GSEA pathway analysis was performed on the 24 h (E,E)- bisantrene treated RNA-seq and proteomic datasets (**Figure 4A, Table S5**). Several commonalities were identified between the cell lines, including activation of coagulation, epithelial to mesenchymal transition (EMT), P53 signaling, and inflammation associated pathways in the transcriptomes of MOLM13 and MV4-11, and the proteome of MOLM13 (**Figure 4A**). E2F targets were significantly decreased in both the transcriptome and proteome datasets for MOLM13 and MV4-11. G2M checkpoint, MYC targets, MTORC1 signaling, and cholesterol homeostasis were decreased in the transcriptomes of (E,E)-bisantrene-treated MOLM13 and MV4-11 (**Figure 4A**).

**Figure 4:**
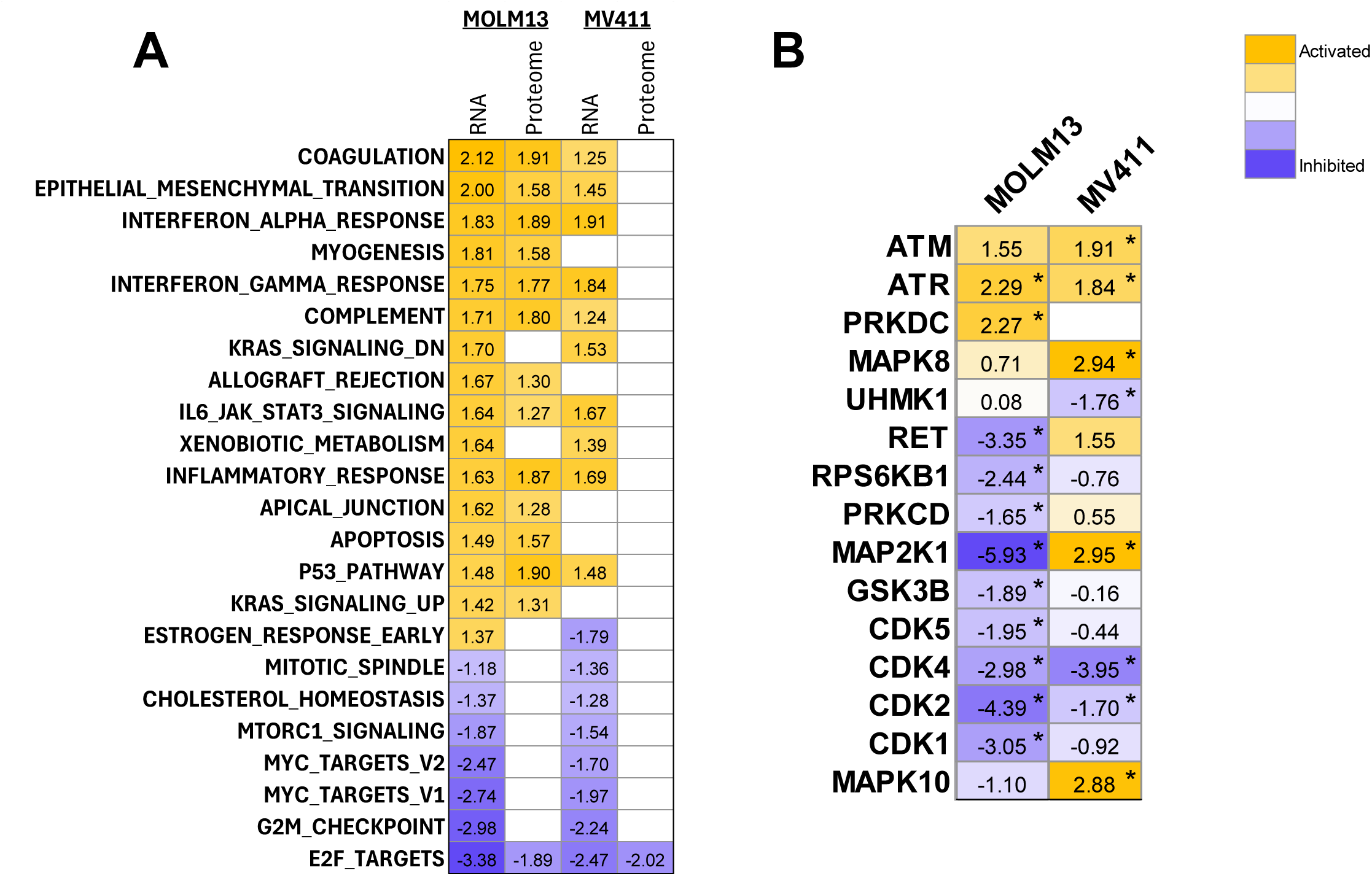
(E,E)-bisantrene treatment reduces MYC and CDK signaling; and increases DNA repair and inflammatory pathways. **(A)** Gene set enrichment analysis of RNAseq and proteomics datasets for MOLM13 and MV4-11 cells treated with (E,E)-bisantrene for 24 h. Heatmap of normalized enrichment scores, where a positive score indicates positive enrichment. Only results with FDR q value <0.25 are shown. Hallmarks with significant enrichment in at least 2 datasets are shown, full result lists are in Table S5 **(B)** Kinase enrichment analysis of activated and inhibited kinases in response to (E,E)-bisantrene inferred from phosphoproteomic data. Positive scores indicate predicted activation; negative scores indicate predicted inhibition. Asterisks indicate significant (p<0.05) results.

### 3.7 Phosphoproteomic analysis reveals activation of DNA repair kinases, and inhibition of CDK activity by (E,E)-bisantrene

To identify protein activity changes in response to (E,E)-bisantrene treatment, phosphoproteomic analysis of MOLM13 and MV4-11 cells treated with (E,E)-bisantrene for 24 h was performed using EasyPhos (**Tables S6** and **S7, Figure S4**). A total of 68 (MOLM13) and 29 (MV4-11) phosphopeptides were significantly increased, and 69 (MOLM13) and 28 (MV4-11) were significantly decreased. In MOLM13, splicing-related factors PRPF4B S366/S368 and SRSF10 S156/S158/S160, Serine/threonine-protein kinase 11-interacting protein S387/S389, and Cyclin-dependent kinase inhibitor 1 S130 were among the top significantly increased in MOLM13 (**Table S6**). Ras-related protein Rab-8A S185, Serine/arginine repetitive matrix protein 2 T983, Mediator of DNA damage checkpoint protein 1 S376/T378, and Fos-related antigen 2 S230 were significantly decreased in MOLM13 (**Table S6**). In MV4-11, Protein phosphatase 1 regulatory subunit 7 S12, Protein PML S527/S530, Protein phosphatase Mg2+/Mn2+ dependent 1G T177, and Unconventional myosin-IXb S1331 were significantly increased, and Prostaglandin E synthase S148/S151, Serine/threonine-protein kinase WNK1 S2011/S2012, Formin-binding protein 1 S359, La-related protein 1 S90, and Ubiquitin-associated protein 2-like S116 were significantly decreased, along with RB1 at multiple sites (S249, S608, S612, S307, S811) (**Table S7**).

There were few overlapping changes between the cell lines, with 5 phosphopeptides commonly increased (Structural maintenance of chromosomes protein SMC1A S957, mitogen activated protein kinase MAPK14 S2, DNA replication complex protein GINS2 S182, Eukaryotic translation initiation factor subunit EIF3K S217, and U3 small nucleolar RNA-associated protein 14 homolog UTP14A S445), and 2 were commonly decreased (Retinoblastoma-like protein RBL1 S640, and G1/S-specific cyclin-E2 CCNE2 S21; (**Tables S6** and **S7**)).

To infer kinase activity changes in response to (E,E)-bisantrene, kinase enrichment analysis was performed using KSEApp (**Figure 4B, Table S8**).^44^ DNA repair kinases ATM and ATR showed activation in MOLM13 and/or MV4-11 in response to (E,E)-bisantrene (**Figure 4B**). Activity of cyclin dependent kinases CDK1, CDK2, CDK4 and CDK5 was significantly reduced in MOLM13 cells. Significant inhibition of CDK2 and CDK4 was also observed in MV4-11 cells. In a complementary approach, kinase enrichment analysis was also performed using PTM-SEA (**Figure S5, Table S9**). Consistent with the results from KSEApp, activation of ATM and ATR and inhibition of CDK1/2/6 was identified in MOLM13 cells. Inhibition of CDK2/6 was identified in MV4-11 cells (**Figure S5, Table S9**).

### 3.8 Regulator analysis reveals inhibition of MYC transcriptional activity and activation of p53 in response to (E,E)-bisantrene

To identify potential drivers of the gene and protein expression changes in response to (E,E)- bisantrene, Ingenuity Pathway Analysis software was used to identify upstream regulators. Tumor suppressor TP53 was the top activated molecule identified in the RNA and proteome datasets of both MOLM13 and MV4-11 cells (**Figure 5A, Table S10**). Other molecules with increased activity included tight junction protein MAGI1, interleukin IL1B, and tumor suppressors CDKN2A and RB1. Peroxisome proliferator-activated receptor PPARD was identified with the greatest reduction in activity in MOLM13 and MV4-11 transcriptomes and concordantly showed decreased activity in the MOLM13 and MV4-11 proteomes (**Figure 5A**). Proto-oncogene MYC displayed the next greatest reduction in transcriptomic activity, with a significant reduction in activity also identified in the MOLM13 proteome (**Figure 5A**). Other molecules with decreased activity included Rab-like protein RABL6, transcription factor CEBPB, growth factor receptor ERBB2, and transcription factor family E2F (**Figure 5A, Table S10**).

**Figure 5:**
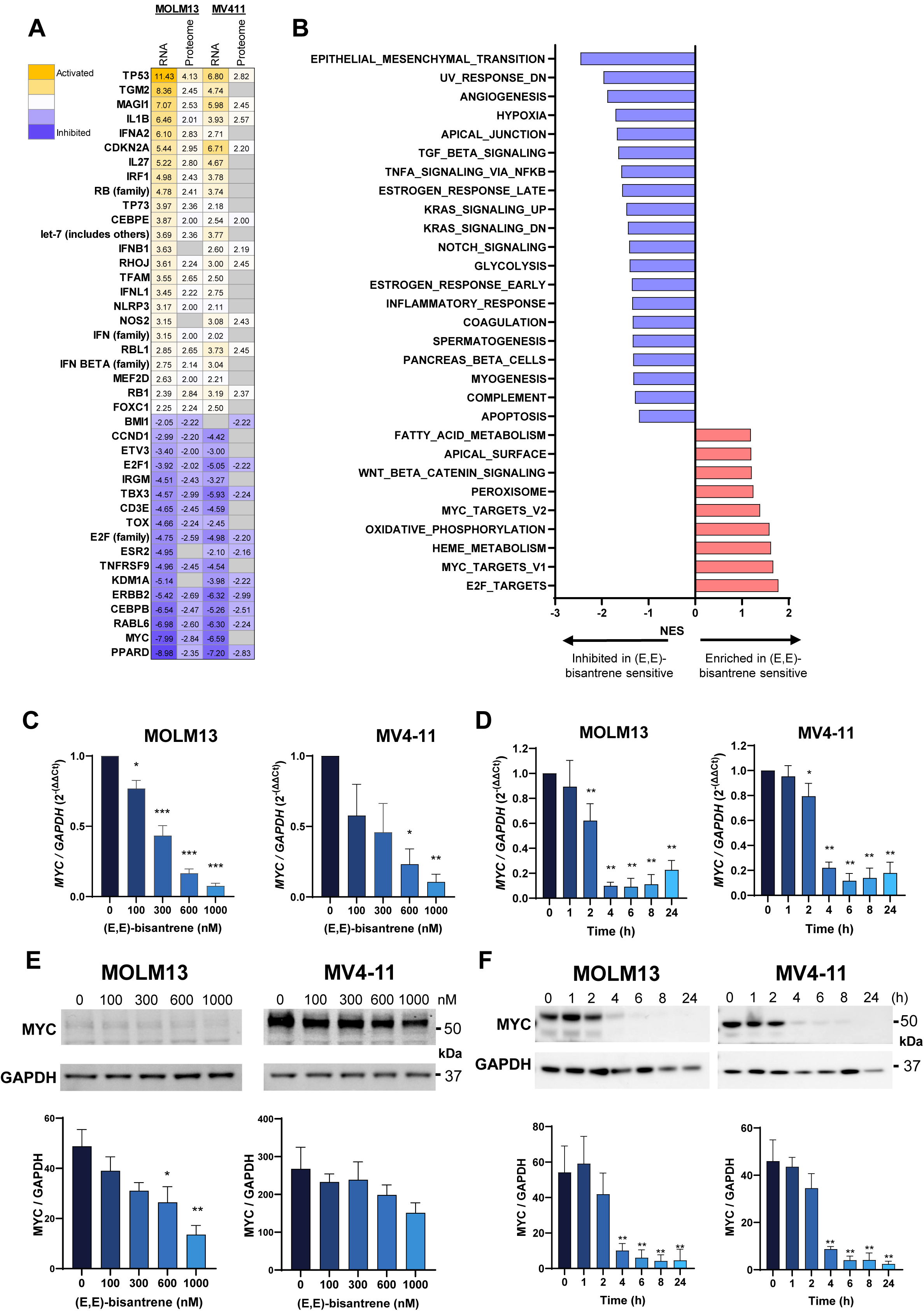
Upstream regulator analysis reveals reduced MYC and increased P53 activity in response to (E,E)-bisantrene. **(A)** Ingenuity Pathway Analysis software was used to identify upstream regulators in RNAseq and proteomics datasets for MOLM13 and MV4-11 cells treated with (E,E)-bisantrene for 24 h. Heatmap of z (activity) scores are shown (p<0.05), where a positive score indicates a predicted increase in activity and vice versa. Results were filtered for entries with p<0.05 in 3 or more datasets, the full results list is provided in Table S10. **(B)** GSEA of human AML cell lines. Transcriptomes of M07e, MV4-11, MOLM13, U937 ((E,E)-bisantrene sensitive) and THP1, OCI-AML3, MonoMac6, HL60, and Kasumi1 (lower sensitivity to (E,E)-bisantrene) were downloaded from DepMap and analysed by GSEA. Hallmark pathways significantly enriched or inhibited in (E,E)-bisantrene sensitive cells are shown. NES=normalized enrichment score. **(C)** *MYC* transcript levels were quantified by qPCR in MOLM13 and MV4-11 following 2 h (E,E)-bisantrene treatment. **(D)** *MYC* transcript levels were quantified by qPCR in MOLM13 and MV4-11 following 300 nM (E,E)-bisantrene treatment at indicated time periods. **(E)** MYC protein levels were evaluated by immunoblot in MOLM13 and MV4-11 following 2 h (E,E)-bisantrene treatment. **(F)** MYC protein levels were evaluated by immunoblot in MOLM13 and MV4-11 following 300 nM (E,E)-bisantrene treatment for indicated time periods. *p<0.05, **p<0.01, ANOVA. Data normalized to GAPDH is shown.

Upregulation of MYC and loss of function of TP53 are both common events in cancer, including AML. MYC has been found to be overexpressed in up to 90% of AMLs^11^ and gene alterations of *TP53* occur in 13% of AMLs.^50^ There is crosstalk between the two transcription factors and they can regulate each other.^51, 52^ MYC is an attractive cancer drug target, however therapeutic inhibition of MYC has proven challenging.^53^ Analysis of transcriptomes^41^ of cell lines more sensitive to (E,E)-bisantrene (i.e., MV4-11, MOLM13, U937, and M07e (**Figure 1A**) compared to cell lines with lower (E,E)-bisantrene sensitivity (i.e., OCI-AML3, THP1, MonoMac6, HL60, and Kasumi1) revealed increased E2F and MYC signatures, and decreased EMT signatures in cell lines more sensitive to (E,E)-bisantrene (**Figure 5B**). Targeted qPCR and western blot analysis of *MYC* transcript and protein levels in MV4-11 and MOLM13 in response to 100, 300, 600, and 1000 nM (E,E)-bisantrene revealed a concentration- and time-dependent decrease. After 2 h treatment with (E,E)-bisantrene, a concentration-dependent reduction in *MYC* transcript levels was observed (**Figure 5C**). A similar trend was observed at the protein level; however, this did not reach significance in MV4-11 cells (**Figure 5E**). Time course analysis with 300 nM (E,E)-bisantrene identified that maximal reduction in *MYC* gene expression occurred by the 4 h time point (**Figure 5D**). A similar trend was observed at the protein level, where almost complete loss of MYC protein expression was observed after 6-8 h treatment with (E,E)-bisantrene (**Figure 5F**). Mutation status of *TP53* or *MYC* was not associated with level of (E,E)-bisantrene sensitivity in the cell lines studied (**Table S1**).

We next analyzed the direct interactors of MYC and TP53 to potentially identify critical effector genes. Protein-protein interaction analysis was performed for the MOLM13 and MV4-11 (E,E)-bisantrene treated transcriptomic datasets (**Figure 6A,B, Figures S6-7**). This identified transcription factor E2F1, homology directed DNA repair protein BRCA1, and transcription regulator CDCA7L as MYC interactors downregulated in response to (E,E)-bisantrene, in both cell lines. E2F1 drives cell cycle progression, suggesting that inhibition of E2F1 may contribute to the G1 accumulation observed in response to (E,E)-bisantrene (**Figure 1C**). TP53 negative regulator MDM2, and cyclin dependent kinase inhibitor CDKN1A were identified as TP53 interactors upregulated in response to (E,E)-bisantrene, in both cell lines (**Figure 6B, Figures S6-8**). Therefore, CDKN1A activation may also contribute to the cell cycle alterations observed in response to (E,E)-bisantrene (**Figure 1C**). Western blotting confirmed a dose-dependent induction of p53 protein, and concomitant induction of p21 (CDKN1A) levels, in response to 2 h treatment with (E,E)-bisantrene (**Figure 6C,D**).

**Figure 6.**
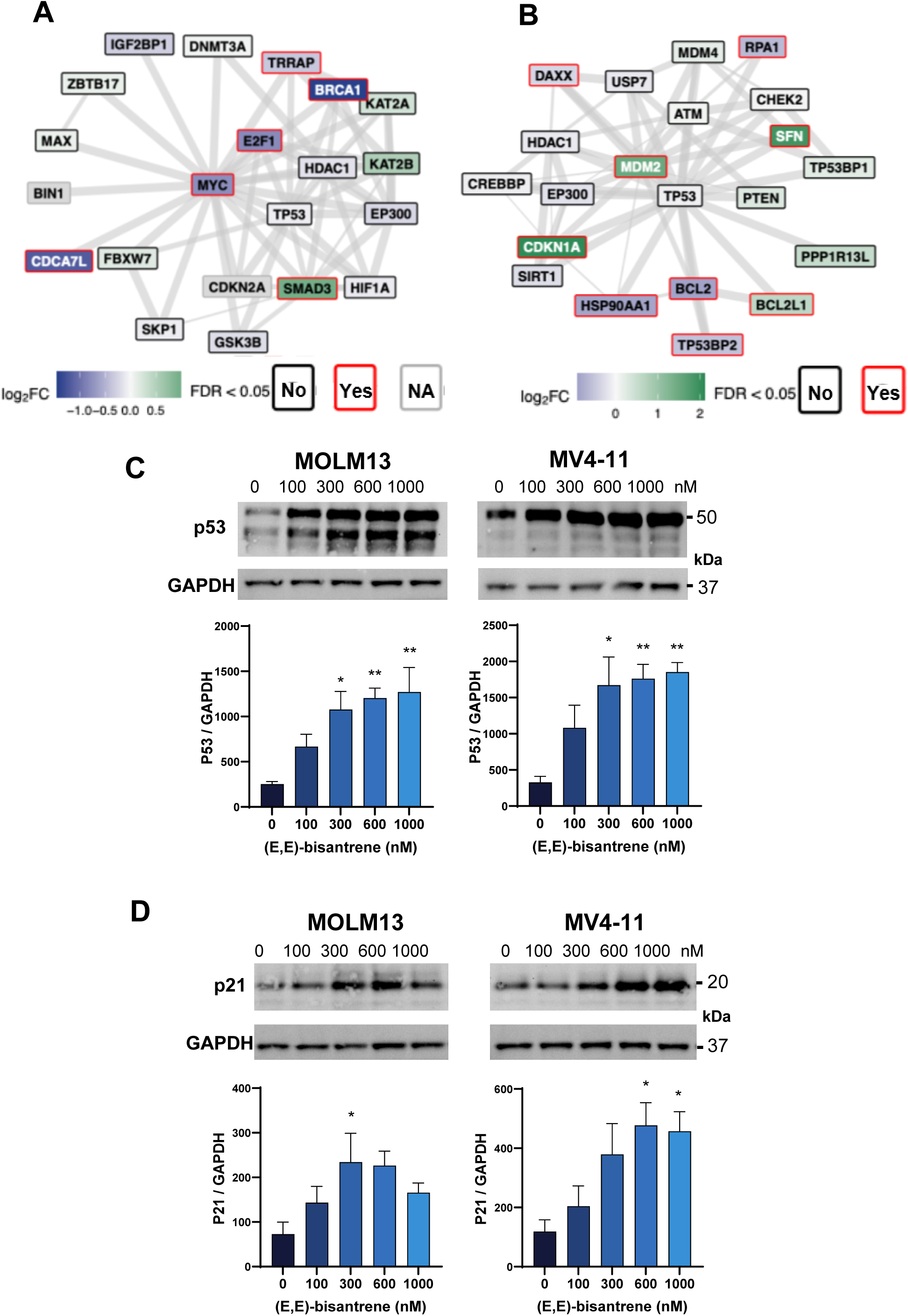
(E,E)-bisantrene induces activation of p53 and p21. **(A,B)** Protein-protein interaction network of (A) MYC and (B) TP53 for MOLM13 cells, generated using STRINGdb revealed interaction with TP53 and CDKN1A (p21). The top 20 nodes ranked by interaction confidence (minimum confidence score ≥ 0.9) and supported by both experimental and curated database evidence were included. Node colour denotes mRNA expression changes in response to (E,E)-bisantrene (blue = downregulated, green = upregulated). Node outlines depict statistical significance from the differential expression analysis (red = *FDR* < 0.05, black = *FDR* ≥ 0.05). Edge thickness corresponds to the STRINGdb interaction confidence scores. (**C,D)** P53 and p21 protein levels respectively were evaluated by immunoblot in MOLM13 and MV4-11 following 2 h (E,E)-bisantrene treatment. *p<0.05, **p<0.005, ***p<0.001, ANOVA. Data normalized to GAPDH.

## 4 Discussion

Bisantrene has demonstrated clinical efficacy in AML and was approved as a salvage treatment for R/R AML in France in 1988, but never marketed.^15^ Recently, (E,E)-bisantrene was identified as the pharmacologically active isomer of bisantrene. The active isomer induced a 40% response rate when used as a monoagent salvage therapy in heavily pretreated R/R AML patients (median of 4 prior lines of therapy).^25^ Importantly, bisantrene has been shown to be well tolerated in >1500 patients in nearly 50 historical clinical trials.^15^

(E,E)-bisantrene can bind and stabilize G4 DNA structures in the promoter region of the *MYC* oncogene, leading to suppression of its transcriptional activity.^14^ Since MYC plays a critical role in AML progression and therapeutic resistance,^10, 11^ this study explored the activity of (E,E)- bisantrene in AML models and its mechanisms of action. (E,E)-bisantrene elicits potent activity in a range of AML models (cell lines *in vitro*, human AML mononuclear cells *ex vivo*, and *in vivo* CDX and PDX models). Mechanism of action studies profiling gene expression, protein expression and phosphoproteomic changes identified suppression of *MYC* expression consistently across different analyses and AML cell lines. Activation of tumor suppressor TP53 signaling and induction of DNA repair and inflammation associated signaling were also observed.

The majority of AML cell lines tested were highly sensitive to (E,E)-bisantrene irrespective of the original driver mutation(s). The most sensitive cell lines were M07e, and *FLT3-ITD* positive (MOLM13 and MV4-11), whereas the *KIT* mutant Kasumi1 was the least sensitive (**Figure 1A**). To analyze if the mutation sensitive associations were causative, isogenic FDPC1 lines transduced with *FLT3*, *KRAS*, or *KIT* mutations, were studied. No differences were observed in (E,E)- bisantrene sensitivity between the isogenic lines (**Figure S1**), suggesting the lower sensitivity of (E,E)-bisantrene in Kasumi1 may be driven by other mechanisms. It should be noted that the mutations in the *KIT-D816V* and Kasumi1 cell lines are not identical and further investigations are required to conclusively demonstrate that oncogenic mutant *KIT* does not induce (E,E)- bisantrene resistance. Notably, all cell lines that showed poor sensitivity to (E,E)-bisantrene had higher basal expression of the multi-drug efflux pump, P-gp. (E,E)-bisantrene is a well-established P-gp substrate,^48^ which may explain the decreased sensitivity of cell lines with high levels of P- gp expression.

To assess the clinical relevance of our results, AML patient derived mononuclear cells were tested for sensitivity to (E,E)-bisantrene *ex vivo*. This analysis identified a range of sensitivities, with *NPM1* mutant samples notably displaying high sensitivity to (E,E)-bisantrene (**Figure 2**). Interestingly, (E,E)-bisantrene was found to increase NPM1 phosphorylation (**Figure S4, Tables S6 and S7**), which can regulate NPM1 cellular localization.^54, 55^ Further studies to assess whether cellular localization of NPM1 affects the (E,E)-bisantrene response is warranted.

(E,E)-bisantrene treatment resulted in a dose-dependent inhibition of leukemia growth and increase in survival in *FLT3-ITD* positive CDX and *FLT3-ITD/NPM1* mutant PDX mouse models (**Figure 3**). Previous work by Su et al.^13^ reported *in vivo* activity of bisantrene in an AML PDX with both *FLT3-ITD* and *FLT3-D835H* mutations, as well as PDXs carrying *MLL-AF9* and *NRAS*- mutations, or *TP53* and *CBFB* deletion. Together, these results suggest that a range of AML mutation subgroups may benefit from (E,E)-bisantrene therapy.

To further investigate the mechanism of action of (E,E)-bisantrene in AML, we performed transcriptomic, proteomic, and phosphoproteomic analysis of MOLM13 and MV4-11 cells treated with (E,E)-bisantrene (**Figure 4**). This identified decreased activity of E2F and MYC targets, as well as decreased activity of a number of cyclin dependent kinases (CDKs, **Figure 4**). Reduced E2F and CDK activity may align with G1 accumulation following (E,E)-bisantrene treatment. Concurrently, (E,E)-bisantrene treatment activated TP53 signaling, EMT, and inflammation associated pathways. Consistent with these results, western blot analysis revealed a dose dependent induction of p53, and induction of p53 target p21 (CDKN1A) levels in response to (E,E)-bisantrene (**Figure 6**). The phosphoproteome revealed activation of DNA double strand break repair signaling (**Figure 4**). As (E,E)-bisantrene is known to inhibit topoisomerase II,^21^ activation of the DNA damage repair response is expected. While DNA double strand break repair signatures were activated, several homology-directed repair signaling transcripts were down- regulated, possibly suggesting a reliance on non-homologous end joining double strand break repair. As homology-directed repair is mostly active in S and G2 phases, while cells preferentially use non-homologous end joining during G1 phase,^56^ this result is consistent with the G1 accumulation in response to (E,E)-bisantrene (**Figure 1**).

Upstream regulator analysis identified TP53 as one of the highest activated, and MYC as one of the most inhibited transcriptional regulators in response to (E,E)-bisantrene exposure (**Figure 5**). Suppression of *MYC* expression by bisantrene has been previously reported.^13^ Here, qPCR and western blot analysis confirmed a reduction of *MYC* transcript and protein levels in MV4-11 and MOLM13 cell lines, in response to (E,E)-bisantrene (**Figure 5**). Notably, GSEA analysis showed that cell lines sensitive to (E,E)-bisantrene had activation of MYC-target gene pathways, suggesting that activated MYC signaling could potentially serve as biomarker for selection of patients likely to respond to (E,E)-bisantrene treatment.

MYC has been shown to be overexpressed in >90% of AML patients, with high levels of MYC leading to poor prognostic outcomes.^11^ Previous studies have emphasized the promise of targeting MYC in AML,^57^ but its intrinsically disordered protein structure has proven to be a notoriously difficult target to modulate using conventional small molecule inhibitors. Our multi-omics data consistently showed that (E,E)-bisantrene could potently suppress MYC and associated molecules (e.g., E2F^58^ and CCND1^59^) in both transcriptomic and proteomic analysis. (E,E)- bisantrene was recently discovered to bind the G4 DNA structure in the promoter region of *MYC* gene, resulting in transcriptional suppression.^14^ This mechanism enables (E,E)-bisantrene to inhibit the activity of *MYC* without directly binding to the MYC protein.

While these multi-omics studies were performed using AML cell lines, the commonality of these oncogenic signaling pathways across multiple cancers suggests therapeutic opportunities are not limited to AML. Indeed, bisantrene has previously shown efficacy in other solid and liquid cancers with elevated MYC expression, including breast cancer.^15^

The pathways targeted by (E,E)-bisantrene suggest intriguing combinations with other AML therapies. Increased expression of HLA transcripts (**Figure S2**), combined with activation of inflammation associated pathways (**Figure 4**), supports exploring the potential utility of (E,E)- bisantrene with immunotherapy. Indeed, (E,E)-bisantrene treatment has been shown to increase the sensitivity of AML cells to T cell killing.^13^ Induction of apoptosis (**Figure 1**) and apoptosis signaling (**Figure 4**) suggests potential combination with the BCL2 family inhibitors. Potentiation of venetoclax activity with (E,E)-bisantrene has been reported.^60^ Many of the (E,E)-bisantrene- targeted pathways identified in this study have previously been linked to resistance to targeted therapies (MYC,^61^ cholesterol biosynthesis,^62^ MTOR^63^). The multiple mechanisms of action of (E,E)-bisantrene may help delay or prevent resistance arising to targeted therapies, particularly in heterogenous multi-clonal cancers like AML. Future preclinical and clinical studies are warranted exploring the potential of (E,E)-bisantrene combination therapies to overcome resistance to targeted agents in both hematological and solid malignancies.

## Availability of data and materials

The datasets used and/or analyzed during the current study are available from the corresponding author on reasonable request. Mass spectrometry data have been uploaded to PRIDE, with the dataset identifier PXD083214 and DOI: 10.6019/PXD083214, and the RNAseq data to the Gene Expression Omnibus, accession number GSE34684.

## Author information

HM, JB, NP, KM, IR, CP, PC and MC conducted cell experiments. HM, JB, KM and AW conducted mouse experiments. HM, JB, NP, KM, IR, MC, SS and NMV conducted data analysis and figure generation. HM conducted proteomics and phosphoproteomics; DK conducted RNA-seq analysis; SS and NMV prepared the original draft of the manuscript, all authors contributed to review. NMV, HM, BJB, SS, MK, DT and MM contributed to project design, interpretation and drafting. NMV provided project management.

## Acknowledgements

The authors thank the Haematology Research and Clinical Trials Unit, Calvary Mater Newcastle Hospital for patient recruitment and sample collection.

## Abbreviations

AML: acute myeloid leukaemia
ATM: ataxia-telangiectasia mutated
BCL2: B-cell lymphoma 2
CDK: cyclin-dependent kinase
DNA-PK: DNA-dependent protein kinase
FLT3-ITD: Fms-like tyrosine kinase 3-internal tandem duplication
IC_50_: half maximal inhibitory concentration
IPA: ingenuity pathway analysis
MS/MS: tandem mass spectrometry
PDX: patient-derived xenograft.

