## Supplementary Table 1 for "(E,E)-bisantrene suppresses *MYC* expression and displays anti-leukemic activity in acute myeloid leukemia"

**Table S1: Anti-proliferative activity of (E,E)-bisantrene in a panel of human AML cell lines.**

| Cell Line Information^ | | | | | | | |
|---|---|---|---|---|---|---|---|
| **Cell Line** | **$IC_{50}$ (nM)*** | **FAB Classification** | **Other Patient Details** | **Translocation/gene fusions** | **Key Mutations** | **CNV (MYC and TP53)** | **P-gp Expression ($log_2$(TPM+1)) **** |
| **NB4** | 25 | M3 | Relapse | PML-RARA | FLT3 E709*, KRAS A18D, TP53 R248Q, FGFR3 S249F | MYC loss | 0.04 |
| **MV4-11** | 26 | M5 | Diagnosis | MLL-AF4 | FLT3-ITD | MYC gain | 0.10 |
| **MOLM13** | 32 | M5a (evolved from MDS) | Relapse | MLL-AF9 | FLT3-ITD, AFF3 T257A, MLH1 V348D, CBL p.?, ZFX Y813fs*4 | MYC amp | 0.02 |
| **AP1060** | 34 | M3 | Relapse | PML-RARA, ETV6-NTRK3 | - | | |
| **HL60** | 126 | M2 | Diagnosis | | NRAS Q61L, CDKN2A pR80*, TNC A39T | TP53 deletion, MYC amp | 0.04 |
| **PL21** | 137 | - | | | KRAS A146V, LATS2 p.?, DNM2 R770*, POLQ V310G, MYC P74L, RPL22 K16fs*9, TP53 P36fs*8, PTMA p.?, ASXL1 I597fs*106, EZH2 D730fs*1 | | 0.04 |
| **NOMO1** | 188 | M5a | Relapse | MLL-AF9 | KRAS G13D, ASXL1 R693*, POLQ V310G, HLA-B E69G, CDKN1B P191fs*35, TP53 C242fs*5, CBLC Q419fs*81, SDHA L649fs*4, PPP6C P52fs*15 | MYC gain | 0.14 |
| **THP1** | 189 | M5 | Relapse | MLL-AF9 | NRAS G12D TP53 R174fs*3, SDHA L649fs*4 | TP53 loss, MYC gain | 0.49 |
| **K-562** | 190 | - | CML | BCR-ABL | TP53 Q136fs*13, AKT3 G37*, PRKCB G294E, EPAS1 Q335*, ASXL1 Y591* | TP53 loss | 0.46 |
| **Kasumi1** | 206 | M2 | Relapse | AML1-ETO | KIT N822K, TP53 R248Q, BCL7A S161W, EPHA7 M893R, CREBBP p.?, SMARCA4 G1370fs*14, NCOA1 p.?, ASXL1 G646fs*12 | TP53 loss, MYC gain | 2.65 |
| **SET2** | 758 | - | essential thrombocythemia | | JAK2 V617F, DNMT3A R882H, SDHA L649fs*4. TP53 R248W, TP53 p.? | | 4.89 |
| **HEL** | 1570 | M6 | Relapse | | JAK2 V617F, TP53 M133K, CM3AP P1267fs*3 | TP53 loss | 5.90 |
| **KG1** | 2514 | | erythroleukemia | | CBLC Q419fs*81, SDHA L649fs*4, TP53 p.?, FOXP1 P584S, HLA-A V100E, ARHGEF10 p.?, NBN p.? | TP53 loss, MYC amp | 5.49 |

| | | | | | | | |
|---|---|---|---|---|---|---|---|
| **OCI-AML3** | - | M4 | Diagnosis | | DNMT3A R882C, NRAS Q61L, NPM1 W288fs*12, BAX E41fs*33, FAT4 p.?, DCC p.? | MYC gain | 0.03 |
| **MO7e** | - | M7 | Diagnosis | | NRAS Q61K, BLM S168*, TSC2 V339I, HLA-A Q78R | MYC amp | |

*Determined using an ATPlite assay after 72 h drug treatment (Oncolines).

**Information collected from DepMap (26Q1 data set). Cell lines with no data have been left blank.

^ Information collected from the DSMZ (https://www.dsmz.de/) and Cell Model Passports[1]

FAB, French-American-British classification. CNV, copy number variation. Amp, amplification. Del, deletion. Fs, frameshift. TPM, Transcripts per million.

1 van der Meer D, Barthorpe S, Yang W et al. Cell Model Passports-a hub for clinical, genetic and functional datasets of preclinical cancer models. Nucleic Acids Res 2019; 47 (D1): D923-d929.
